# *Wolbachia* induce cytoplasmic incompatibility and affect mate preference in *Habrobracon hebetor* to increase the chance of its transmission to the next generation

**DOI:** 10.1101/505073

**Authors:** Zeynab Bagheri, Ali Asghar Talebi, Sassan Asgari, Mohammad Mehrabadi

## Abstract

*Wolbachia* are common intracellular bacteria that are generally found in arthropods including a high proportion of insects and also some nematodes. This intracellular symbiont can affect sex ratio with a variety of reproductive anomalies in the host, including cytoplasmic incompatibility (CI) in haplodiploides. In this study, we questioned if the parasitoid wasp, *Habrobracon hebetor* (Hym.: Braconidae), which is one of the most important biological control agents of many lepidopteran larvae, is infected with *Wolbachia*. To test this, DNA was extracted from adult insects and subjected to PCR using specific primers to *Wolbachia* target genes. The results showed high rate of *Wolbachia* infection in this parasitoid wasp. To find out the biological function of *Wolbachia* in *H. hebetor*, we removed this bacterium from the wasps using antibiotic treatment (cured wasps). Results of the crossing experiments revealed that *Wolbachia*induced CI in *H. hebetor* in which cured females crossed with infected males produced only males, while in the progeny of other crosses, both males and females were observed. Also, our result showed that the presence of *Wolbachia* in the females increased fecundity and female offspring of this parasitoid wasp. However, the presence of *Wolbachia* in the males had no significant effect on the fecundity and female production, but might have incurred costs. We also investigated the effect of *Wolbachia* on mate choice and found that *Wolbachia* affects mating behavior of *H. hebetor*. Together, we show that *Wolbachia* induce CI in *H. hebetor* and affect host mating behavior in favor of its transmission. *Wolbachia* utilize these strategies to increase the frequency of infected females in the host population.

## 1. Introduction

*Wolbachia* are endosymbiotic bacteria that infect more than 52% of arthropod species and also some nematodes. This intracellular symbiont is among the facultative endosymbionts and can be transferred to the next generation by vertical transmission. Utilizing different strategies to expand host rang is an exceptional feature of *Wolbachia*; altering sex ratio with a variety of reproductive anomalies in the host (LePage and Bordenstein, 2013), protection from replication of a variety of pathogens (Hedges et al., 2008; Pan et al., 2017; Yixin et al., 2013), alteration in behavior (Miller et al., 2010; Peng et al., 2008; Rohrscheib et al., 2015; Vala et al., 2004) and some metabolic pathway (Brownlie et al., 2009; Hosokawa et al., 2010) are some of *Wolbachia*’ effects on the host.

*Wolbachia* can alter reproductive system via male-killing (Hurst et al., 1999; Jiggins et al., 1998), feminization in isopods (Rousset et al., 1992), parthenogenesis induction (PI) in haplodiploid species in which unfertilized eggs become females (Stouthamer et al., 1990) and importantly cytoplasmic incompatibility (CI) (Yen and Barr, 1971). CI is commonly expressed when *Wolbachia*-infected males mate with uninfected females (Unidirectional CI) and also occurs in mattings between infected individuals harboring different strains of *Wolbachia* (Bidirectional CI) (Hunter et al., 2003; O’Neill and Karr, 1990; Perrot-Minnot et al., 1996b). In unidirectional CI crosses, paternal chromatin fails to condense properly in the first cell cycle and line on the metaphase plate during the first mitosis. Such a cross results in the death of the embryo (Beckmann et al., 2017). This strategy can be used in the mass production and release of incompatible male insects to control wild populations of disease vectors such as the mosquito *Culex pipiens* (Laven, 1967) and of agricultural pests such as *Liriomyza trifolii* (Tagami et al., 2006).

It has been reported that *Wolbachia* may have positive effects on the fitness of host. For example, it was shown that *Wolbachia* increased fecundity in the parasitoid wasp Trichogramma bourarachae (Vavre et al., 1999). It has also been reported that *Wolbachia* infection positively affects the life history and reproductive traits of Callosobruchus chinensis males and females (Okayama et al., 2016). Given that parasitoid wasps play an important role in the biological control of insect pests, the presence of *Wolbachia* can affect their function. Previous studies have reported the effect of *Wolbachia* on reproduction of some parasitoid wasps; for example, PI-*Wolbachia* has been reported in *Trichogramma* sp. (Trichogrammatidae) (Stouthamer and Werren, 1993) and *Encarsia* sp. (Aphelinidae) (Stouthamer and Mak, 2002; Wang et al., 2017; Zchori-Fein et al., 1992). Also, in the parasitoid wasp Asobara tabida (Braconidae), it has been shown that two strains of *Wolbachia* (i.e. wAtab1 and wAtab2) induce CI, but wAtab3 strain is required for the host oogenesis (Dedeine et al., 2005). CI-*Wolbachia* has been reported form other wasps such as *Aphytis melinus* (Aphelinidae) (Vasquez et al., 2011) and *Nasonia vitripennis* (Pteromalidae) (Breeuwer and Werren, 1993).

*Habrobracon hebetor* (Hym.: Braconidae) is one of the most important biological control agents of many lepidopteran larvae, mainly pyralid moths infesting stored products (Ghimire, 2008). In previous studies, the presence of *Wolbachia* in *H. hebetor* has been reported (Kageyama et al., 2010); however, the effect of *Wolbachia* on this parasitoid wasp has not been investigated yet. Therefore, in this study we questioned whether *Wolbachia* affects the reproduction of *H. hebetor*. Our results indicated the presence of *Wolbachia* in the population of *H. hebetor* collected from Karaj, Iran. We show that *Wolbachia* induce CI in *H. hebetor* and affect the wasp mating behavior in favor of *Wolbachia* transmission.

## 2. Materials and Methods

### 2.1. Insects

The original population of the *H. hebetor* was collected from Karaj, Iran. The wasps were reared under the laboratory condition at 25 ± 5 ºC, 60 ± 5% RH and a photoperiod of 16:8 h and fed daily with diluted raw honey (90% honey and 10% water). The female wasps were presented with fifth instar mill moth (*Ephestia kuehniella*) larvae for oviposition.

### 2.2. *Wolbachia* detection

Total DNA was extracted from single individuals of *H. hebetor* (n=100) using the previously described procedure (O’Neill et al., 1992). In order to detect *Wolbachia*, the DNA samples were subjected to PCR by using the specific primers targeting *Wolbachia wsp* gene (Narita et al., 2007). The PCR reactions were conducted under a temperature profile of 95 °C for 5 min, followed by 35 cycles of 95 °C for 30 sec, 56 °C for 30 sec and 72 °C for 1 min, and a final extension at 72 °C for 10 min. The PCR products were subjected to agarose (1%) gel electrophoresis. Moreover, to compare *Wolbachia* density, DNA was extracted from each developmental stage of the wasp including the egg, larva (1st, 2d, 3th and 4th instar), pupa (3-, 5- and 7-days old), and adult (1-, 15-, 20-days-old males; and 1-, 15-, 20-, 25-, 30-days-old females) as described above. Concentrations of the DNA samples were measured using a Epoch instrument (BioTek), and 10 ng from each genomic DNA sample was used for quantitative PCR (qPCR) using SYBR Green Mix without ROX (Ampliqon) with a Mic real time PCR (BMS) under the following conditions: 95 °C for 15 min, followed by 45 cycles of 95 °C for 10 sec, 30 sec at the annealing temperature, and 72 °C for 30 sec, followed by the melting curve (72-95 °C). Specific primers targeting *Wolbachia ftsZ* gene (Kruse et al., 2017) and the insect 18s *rRNA* gene as a reference gene (Karamipour et al., 2016) were used (Table 1). Reactions from three biological replicates were repeated three times.

**Table 1.**
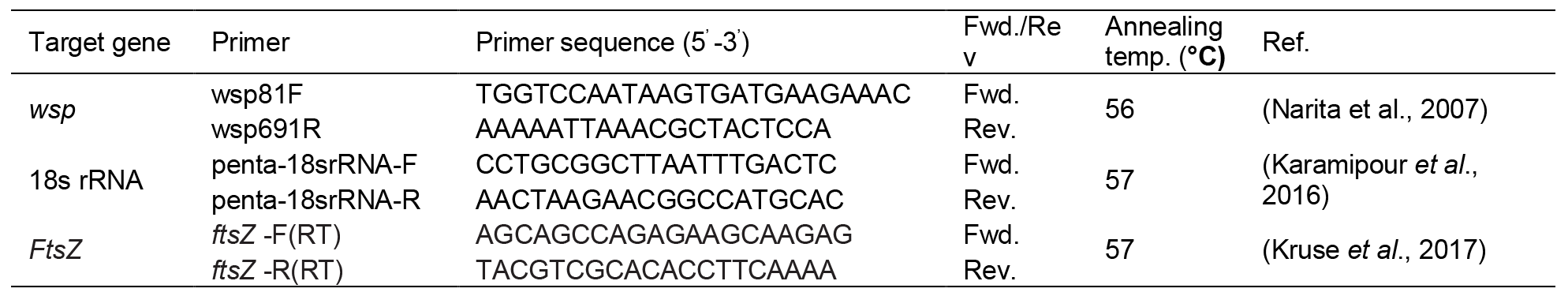
Primers used in this study.

### 2.3. Antibiotic treatments

To find out the biological function of *Wolbachia* in *H. hebetor*, four isolines (i.e. two *Wolbachia*-infected (W^+^) and two uninfected or cured (W^-^)) were generated using tetracycline treatment (0.2%, w/v with diluted raw honey) for seven days. Thereafter, the wasps were allowed to lay eggs on the host, mill moth larvae. Tetracycline treatment was continued for three generations (seven days treatment of young adults per generation), and the number of 10 wasps were then randomly selected to test for *Wolbachia* infection using qPCR as described above. After successful removal of *Wolbachia* from the wasps, they were reared for five generations. The resultant adult wasps from the 8th generation were considered as the uninfected isolines. The infected isolines were also treated the same; however the antibiotic was not used. The adult wasps from these isolines were used for the cross-mating experiments.

### 2.4 Crossing experiments and assessing the role of *Wolbachia* in *H. hebetor*

For the crossing experiments, we collected 100 pupae from the infected and uninfected isolines and allowed them to emerge. Then, 20 pairs of unmated individuals (virgin) from these isolines were used for the crossing tests as follow: infected female—infected male, infected female—uninfected male, uninfected female—infected male and uninfected female—uninfected male. We counted the number of the resultant eggs, rate of hatching eggs, numbers of larvae, pupae and offspring during three days after crossing. We also estimated the sex ratio of the emerged wasps.

### 2.5. Effect of *Wolbachia* on *H. hebetor* mate choice

In order to investigate the effect of *Wolbachia* on mate choice, we placed virgin infected or uninfected females individually in 5-cm-diameter cups with two virgin males, one infected and one uninfected (i.e. a virgin infected female and two virgin males; a virgin uninfected female and two virgin males). To differentiate between the males, either infected or uninfected males (equally) were marked with a point on the end of their wings using a black pen. Forty replicates of each combination were conducted and the rates of successful mating were counted for 15 min. In successful mating, a female accepts a male copulation attempt by moving her ovipositor laterally and then remaining immobile. But a female prevents successful mating by walking, knocking the male with the hind legs, or bending down the abdomen.

### 2.6. Data analysis

*Wolbachia* densities in different developmental stages of *H. hebetor* were compared using ANOVA followed by Tukey’s multiple comparisons test. *Wolbachia* density in the infected (W^+^) and the uninfected (W^-^) parasitoid wasps were compared using unpaired t-test. Comparisons of the mean number of eggs per female wasp within three days in different crosses were analyzed with Two-way ANOVA followed by Tukey’s (HSD) test. The mean number of eggs, the rate of hatching eggs, and the percentage of pupae formation, the percentage of progeny emergence and the sex-ratio of progeny emergence per female in different crosses were compared using Mann-Whitney *U*-test. The percentage of mating was analyzed using Kruskal-Wallis test. Statistical analyses were performed using the software SPSS Statistics 17.0 and the graphs were created by using Graph Pad Prism.

## 3. Results

### 3.1. *Wolbachia* are persistently present during development of *H. hebetor*

To detect and estimate *Wolbachia* infection rate, DNA was extracted from 100 individuals (both sexes) of *H. hebetor*. The results showed that all the tested individuals were infected with *Wolbachia* (Fig. 1A). Also, to estimate the relative density of *Wolbachia* in different developmental stages of *H. hebetor*, DNA samples of each developmental stage (i.e. egg, larva, pupa, and adult) were screened by qPCR. We found that *Wolbachia* was present in all the developmental stages of the parasitoid wasp and its density in the eggs and adults were higher than that of the other stages (Fig. 1B; F_9, 20_ =70.86, P < 0.0001). The density of *Wolbachia* in different larval and pupal stages showed no significant difference. Also, the result showed that *Wolbachia* density increased along with the increase in the age of females (F_4, 10_ = 556.3, P < 0.0001), while *Wolbachia* density did not differ significantly among 1-day, 15-day and 20-day-old males (Fig. 1C).

**Fig. 1.**
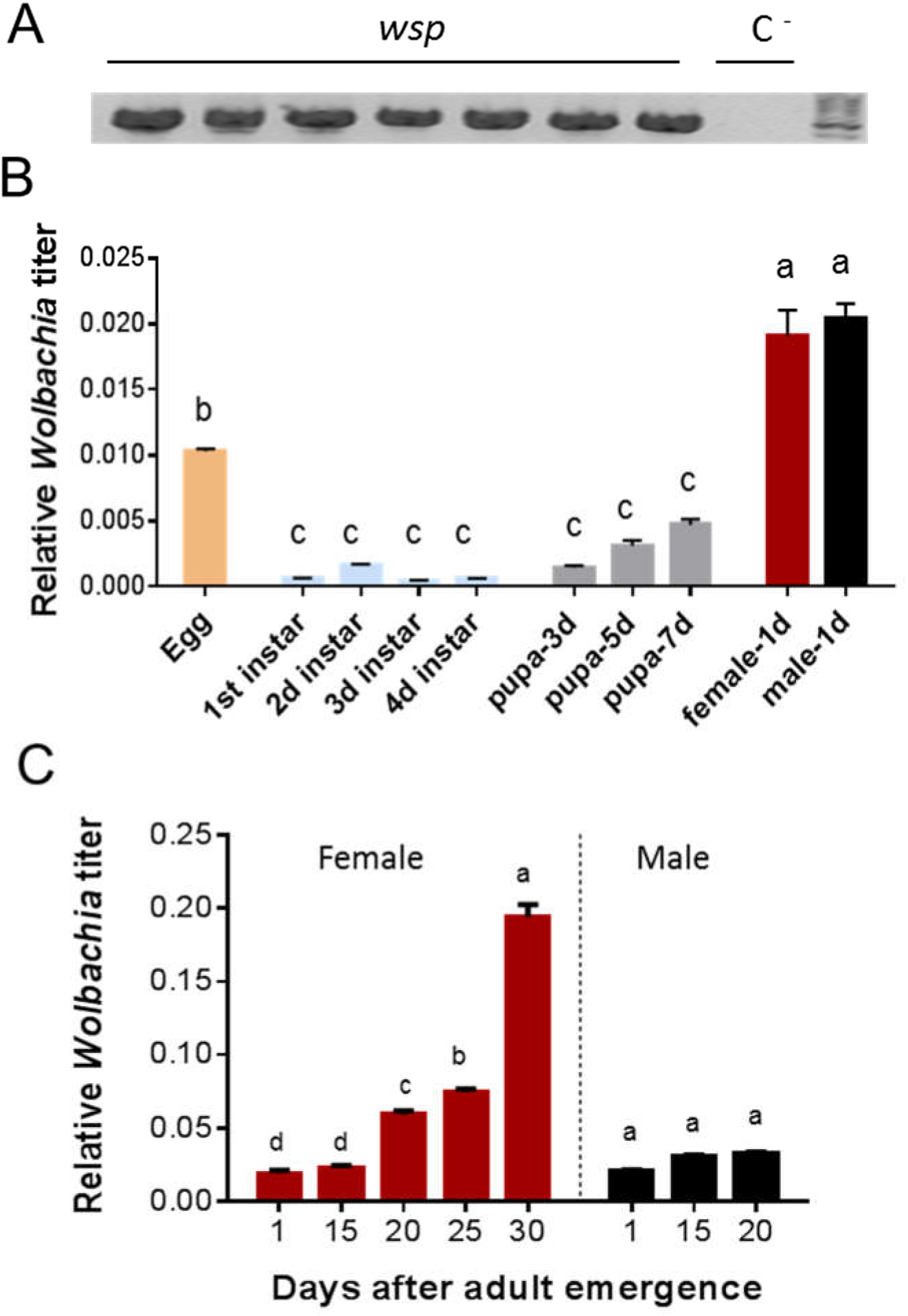
Detection of *Wolbachia* and their density fluctuations during development of H. hebetor. A: A representative diagnostic PCR of *Wolbachia* in adults of *H. hebetor* collected from Karaj using wsp primers. C^-^, negative control lacking DNA sample. B: Quantitative PCR (qPCR) detection of *ftsZ* gene of *Wolbachia* during development of *H. hebetor*. C: Effect of aging on *Wolbachia* density in female and male adult of *H. hebetor*.

### 3.2. *Wolbachia* induce cytoplasmic incompatibility in *H. hebetor*

To investigate the effect of *Wolbachia* infection on the reproduction and fitness of *H. hebetor*, we created uninfected *H. hebetor* isolines by treating the wasps with tetracycline. Following confirmation of removal of *Wolbachia* from tetracycline-treated wasps (Fig. 2A), we tested if the antibiotic treatments affected mitochondria density (Ballard and Melvin, 2007) and microbiota population (Karamipour et al., 2016) in the treated wasps and found no significant differences with the control wasps (Fig. S1). Then, we created four different crosses and analyzed different parameters associated with reproduction of the wasp. The results of the crosses showed that all the crosses produced both males and females progeny, except when infected males were crossed with the uninfected females which resulted in only male progeny (Fig. 2B). It is worth mentioning that in hymenopterans, sex determination is based on haplodiploidy system. In other words, males are haploid and females are diploid. Therefore, CI only appears in the female progeny (Breeuwer and Werren, 1993; Perrot-Minnot et al., 1996a). Our results also suggest that *Wolbachia* induced CI in the parasitoid wasp, *H. hebetor*.

**Fig. 2.**
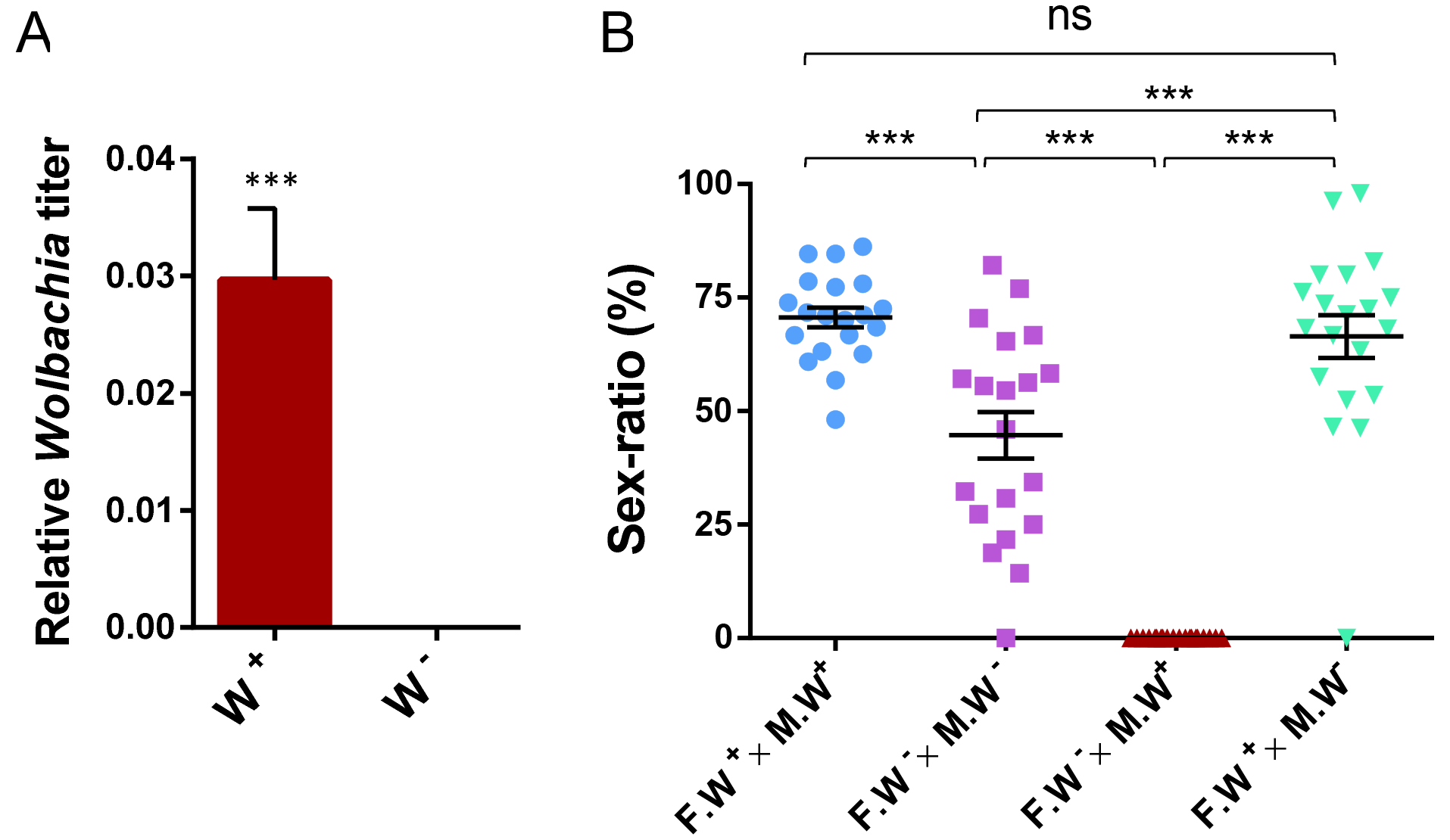
Confirmation of *Wolbachia* removal and the effect of *Wolbachia* on progeny sex ratio. A: Antibiotic treatment resulted in *Wolbachia* elimination in H. hebetor. qPCR analysis of *Wolbachia* density in the *Wolbachia* infected (W^+^) and the *Wolbachia* cured (W^-^) parasitoid wasps. *** P<0.001. B: Sex ratio of the offspring resulted from different crosses during three days oviposition. Each data point represents a total number of female offspring produced per female during three days. Female (F), Male (M), *Wolbachia*-infected (W^+^) and *Wolbachia*-cured (W^-^). Mann-Whitney U test, *** signifies P<0.001.

We also compared the number of resultant eggs of the crosses during three days oviposition. The mean number of eggs within three days per female in infected female-infected male and infected female-uninfected male crosses was significantly higher than those of the other two crosses (Fig. 3A). In addition, the rate of egg hatching in uninfected female-infected male cross was significantly lower than that of other crosses (Fig. 3B). The rates of pupa formation were not significantly different in different crosses (Fig. 3C), while the progeny emergence rate of infected female-uninfected male was significantly higher than that of other crosses (Fig. 3D). Also, the results showed that the presence of *Wolbachia* in both male and female (i.e. infected female—infected male cross) significantly increase the progeny emergence rate compared with the absence of *Wolbachia* (cured female-cured male cross) (Fig. 3D).

**Fig. 3.**
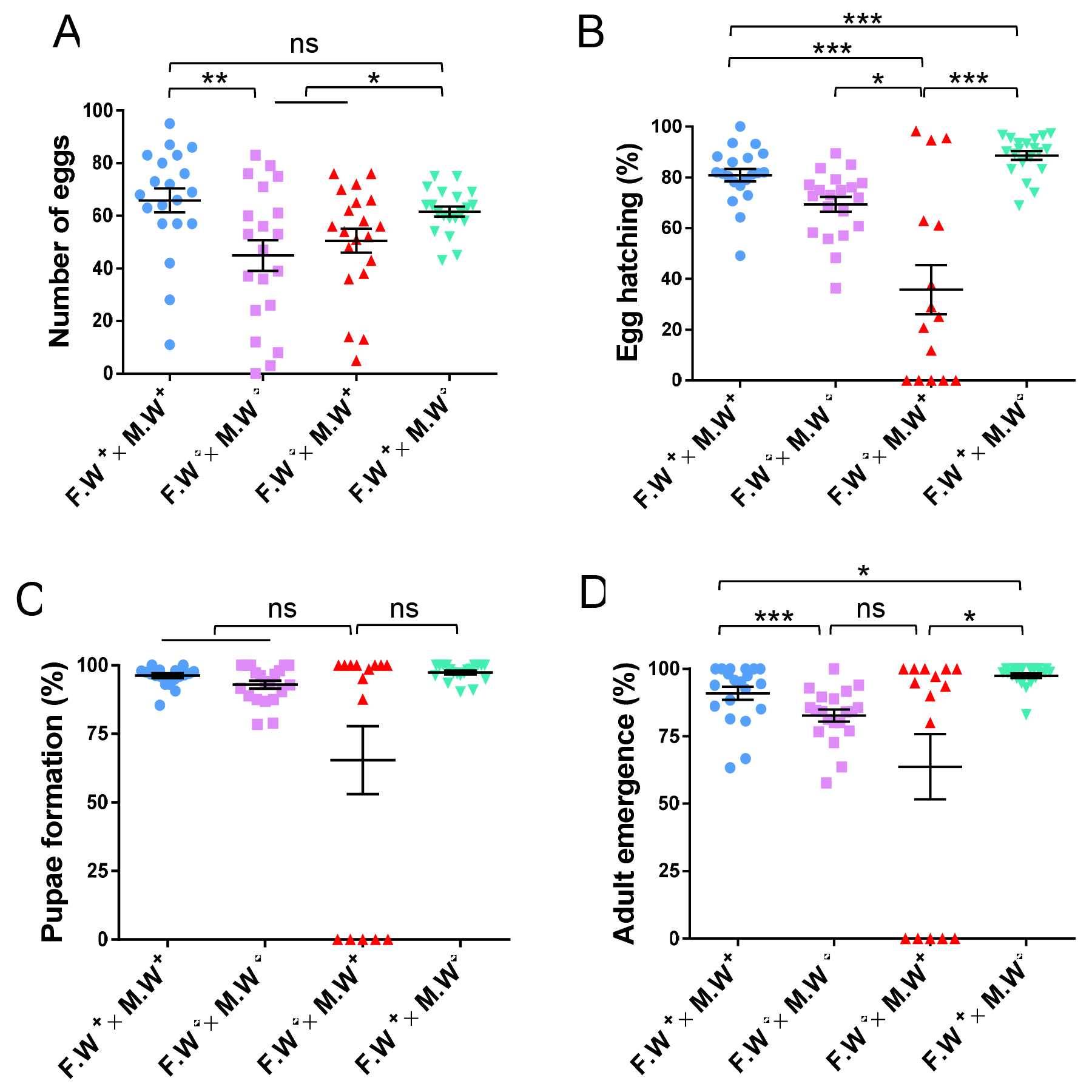
Comparison of biological parameters of *H. hebetor* in different crosses. A: Comparison of the mean number of eggs per female. B: The rate of hatching. C: The percentage of pupae formation. D: The percentage progeny emergence in different crosses. Female (F), Male (M), *Wolbachia*-infected (W^+^) and *Wolbachia*-cured (W^-^). Each data point represents a sum of data obtained per female during three days. Mann-Whitney *U* test, *signifies P < 0.05, **signifies P < P<0.01, *** signifies P<0.001.

Moreover, we compared the mean number of eggs per day in each cross. On the first day, the mean number of eggs in infected female-infected male cross was significantly higher than that of the other crosses. On the second day, there was no significant difference among the crosses in term of the mean number of eggs. On the third day, however, the mean number of eggs in infected female-uninfected male cross was significantly higher than that of others (Fig. 4, Table 2). Notably, the highest numbers of progeny females were observed in infected female—infected male and infected female—uninfected male crosses (Fig. 2B).

**Fig. 4.**
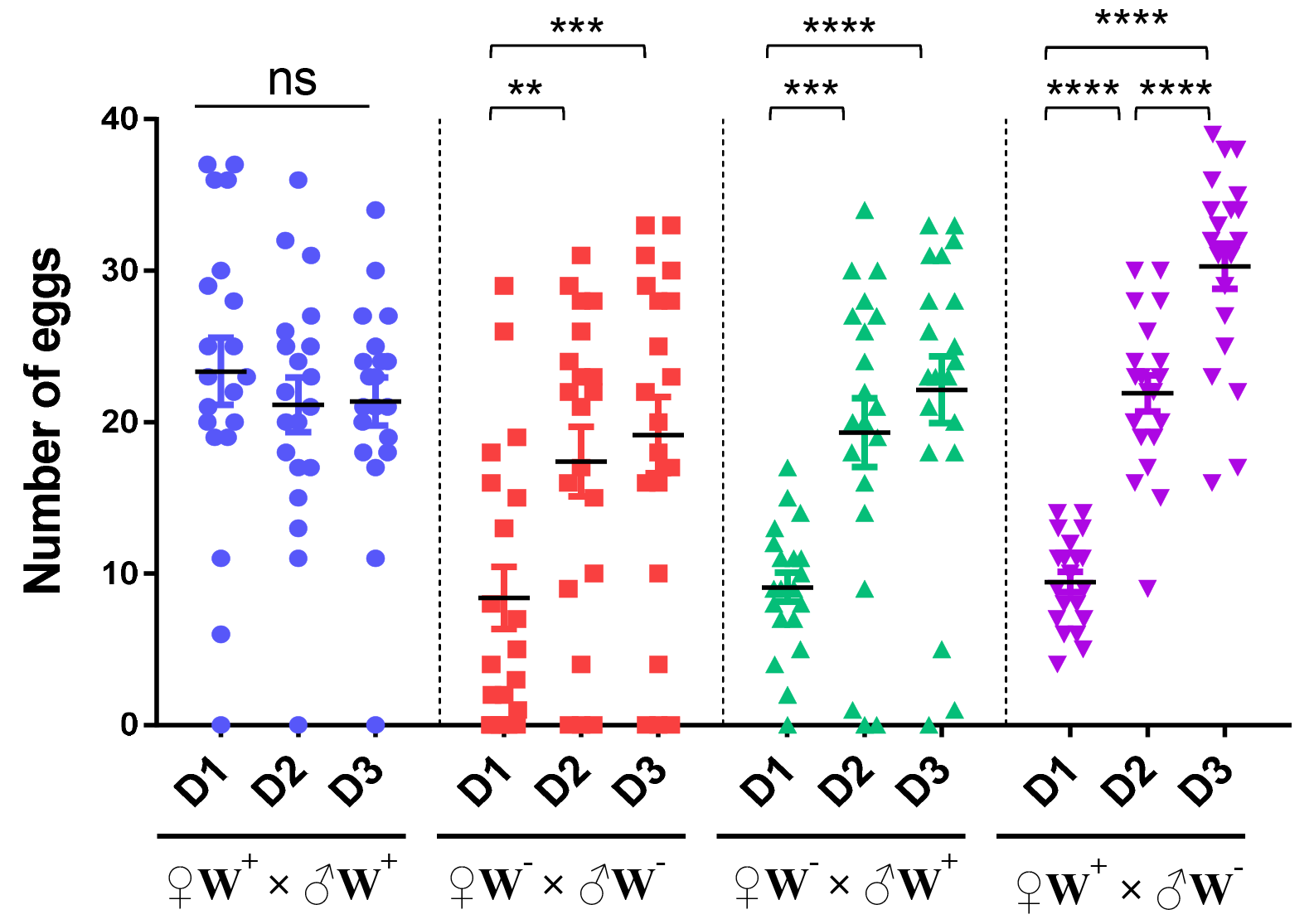
Comparison of the mean number of eggs per female in three days in different crosses. Each data point represents a number of eggs per female. Means were compared by Tukey’s (HSD) test. Day (D), *Wolbachia* infected (W^+^) and *Wolbachia* cured (W^-^). *signifies P < 0.05, **signifies P < 0.01, *** signifies P<0.001, **** signifies P<0.0001

**Table 2.**
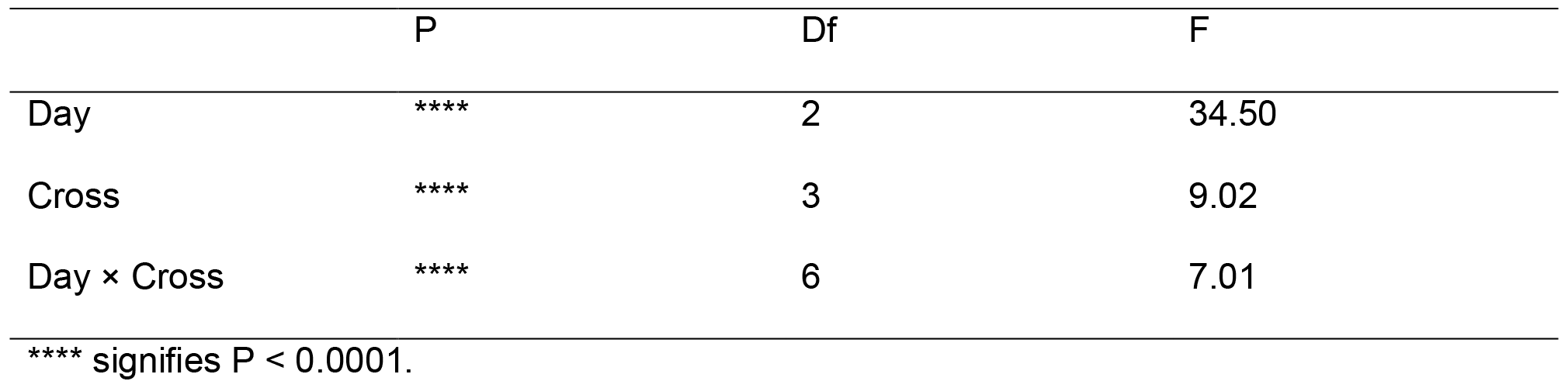
Analysis of Variance (ANOVA). The effect of day on number of eggs in different crosses in *H. hebetor*

|  | P | Df | F |
| --- | --- | --- | --- |
| Day | **** | 2 | 34.50 |
| Cross | **** | 3 | 9.02 |
| Day × Cross | **** | 6 | 7.01 |

\*\*\*\* signifies $P < 0.0001$ .

### 3.3. *Wolbachia* affect mate choice in *H. hebetor*

In order to investigate the effect of *Wolbachia* on mate choice, we designed a choice mating test. In this test, two infected and uninfected males were exposed to either an infected or an uninfected female. The results showed that the infected females mated at a higher rate with infected males compared to the uninfected males (K-W test, x2= 5.00, P = 0.03). However, there was no difference between mate choices of uninfected females (K-W test, x2= 0.8, P = 0.3) (Fig. 5).

**Fig. 5.**
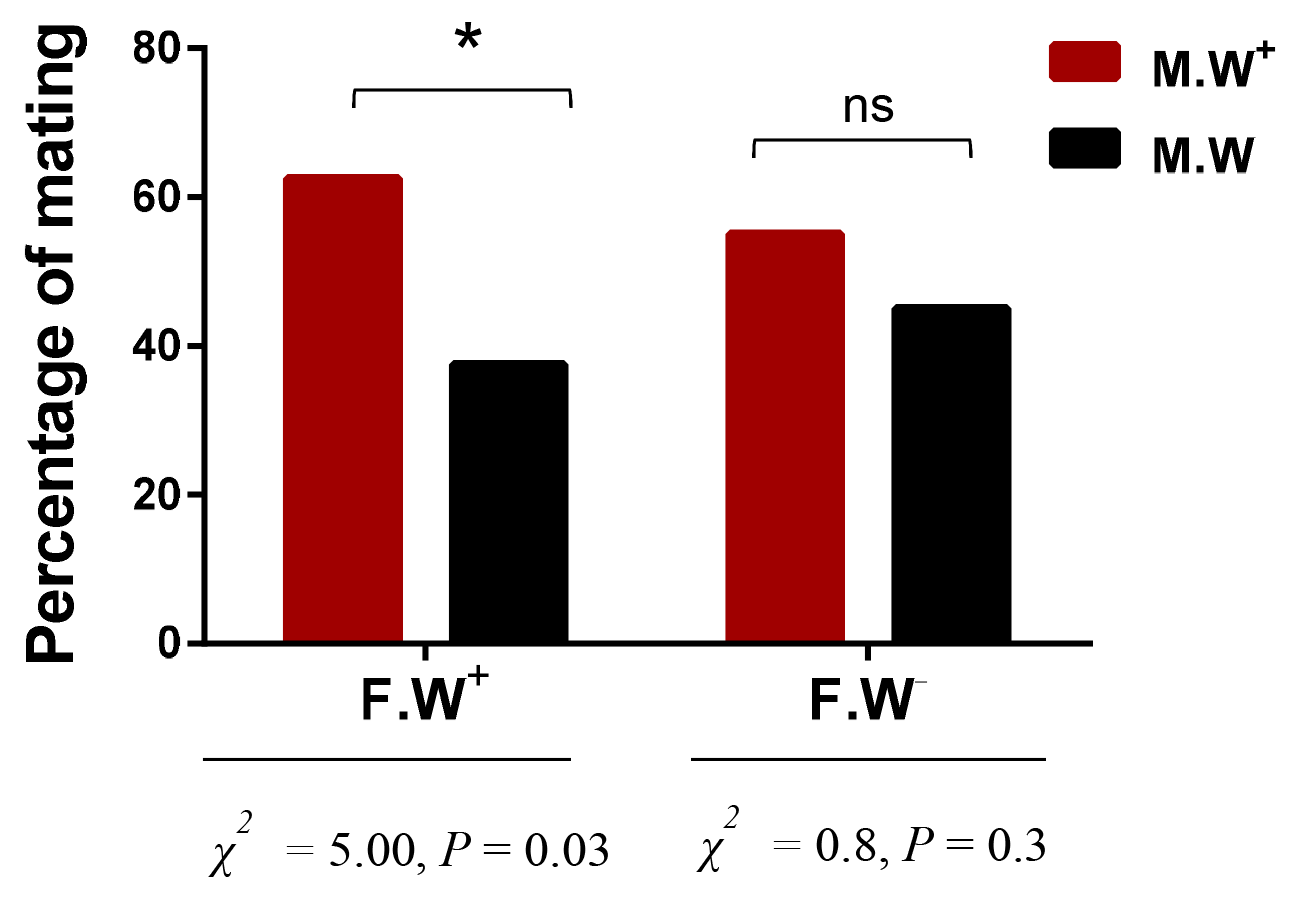
Effect of *Wolbachia* on female mate choice. Female (F), Male (M), *Wolbachia* infected (W^+^) and *Wolbachia*-cured (W^-^). Kruskal-Wallis (K-W) test, * signifies P < 0.05.

## 4. Discussion

*Wolbachia* have evolved several mechanisms of host reproductive manipulations to increase their worldwide prevalence, including feminization, parthenogenesis, male-killing, and cytoplasmic incompatibility (CI). These phenotypes, except CI, serve to increase the frequency of *Wolbachia*-infected females, the transmitting sex of the bacteria, in a host population as males are an evolutionary dead end for *Wolbachia* (Hurst and Frost, 2015; Serbus et al., 2008; Werren et al., 2008). The presence *Wolbachia* has been reported in many insects. Given that *Wolbachia* manipulate reproductive system of their insect hosts, they have good potential in biological control, either to control insect pests or to increase the population of biological control agents and improve their functions.

Here, we investigated the effect of *Wolbachia* on the reproductive biology of the parasitoid wasp *H. hebetor* which is one of the important biological control agents of many lepidopteran larvae. We detected *Wolbachia* from all the field-collected individuals of *H. hebetor* and the prevalence rate of infection was 100%. High rate of *Wolbachia* infection in natural population of *H. hebetor* could be explained by fitness benefit that *Wolbachia* confers to the wasp. Also, we estimated relative density of *Wolbachia* in different developmental stages of *H. hebetor* and found that *Wolbachia* densities in the eggs and adult females were higher than other stages of the parasitoid wasps. These results indicate efficient vertical transmission of *Wolbachia* in this parasitoid wasp. Developmental-specific densities of *Wolbachia* have also been reported in other insects such as *Tribolium confusum* (Ming et al., 2015) and *Brontispa longissima* (Ali et al., 2018) with higher levels in eggs and adults than other stages. *Wolbachia* density in *Diaphorina citri* (Dossi et al., 2014) and Drosophila melanogaster (Goto et al., 2006) was shown to increase as development proceeds. Considering our results and those reported from other studies, it can be hypothesized that *Wolbachia* may affect or be regulated throughout the host development. Moreover, high rate of *Wolbachia*transmission might be achieved by investing high population of the bacteria within the host egg.

In addition, we investigated the effect of ageing on the *Wolbachia* density in the adult males and females. The result showed that *Wolbachia* density increased in aged females that is in congruence to the study that showed *Wolbachia* density increase with aging in Laodelphax striatellus females (Noda et al., 2001). We found no change in *Wolbachia* density over time in the males. Similar to our results, it has been shown that *Wolbachia* density in the males of Tribolium confusum remains unchanged over time (Ming et al., 2015). *Wolbachia* are positively associated with CI strength (Bourtzis et al., 1996; LePage and Bordenstein, 2013; Sinkins et al., 1995). Reduction in *Wolbachia* density during aging in males has been suggested to be associated with reduction in the ability to cause CI. This phenomenon has been reported in *Sogatella furcifera* males (Noda et al., 2001) and Aedes albopictus males infected with *W*AlbA strain of *Wolbachia* (Tortosa et al., 2010). Therefore, based on our results, it could be hypothesized that *Wolbachia*-induced CI increase over time in the female wasps, while it remains almost constant in the males. Males suffer from fertility reduction in CI crosses, therefore by removing/reducing the infection during development they may suppress the expression of CI (Tortosa et al., 2010). In the present study, the density of *Wolbachia* in 1-day, 15-day and 20-day-old males was almost the same, and it can be concluded that males prevent *Wolbachia* growth to suppress CI. However, density of *Wolbachia* increases in females with aging, with 30-day-olds having the highest bacterial density.

Given the high rate of *Wolbachia* infection and their presence during the wasps’ development, we next examined the functional role of *Wolbachia* in this insect. To assess the effect of *Wolbachia* infection on the reproduction of *H. hebetor*, we designed four different crosses and found that the infected males crossed with uninfected females resulted in only male progeny (i.e. incompatible cross). The rates of eggs hatching, pupa formation and adult emergence of this cross were also lower than that of others. Two types of CI have been suggested in haplodiploids: female mortality or conversion to males (Bordenstein et al., 2003). Given low hatching rate and all male progeny in the incompatible cross of our experiments, the mortality type of CI is suggested for *H. hebetor*. Female mortality has been suggested as the normal and most common type of CI in haplodiploid insects including parasitoid wasps such as Nasonia (Perrot-Minnot et al., 1996a), Leptopilina (Vavre et al., 2000) and Trichopria (Vavre et al., 2002), whereas female conversion to males is an exception (Bordenstein et al., 2003).

Our data showed that the presence of *Wolbachia* in the female wasps improved their fecundity. The number of eggs laid by the infected female wasps was significantly higher in the compatible crosses (i.e. infected female-infected male cross and infected female-uninfected male cross) compared with the others. Consistent to these results, it has been reported that *Wolbachia*-infection improves fecundity in *Aedes albopictus* (Dobson et al., 2004), *Trichogramma* (Vavre et al., 1999), *D. melanogaster* (Fry et al., 2004) and Tetranychus urticae (Zhao et al., 2013). Higher fecundity of the infected females would increase infected individuals thereby accelerate *Wolbachia* distribution within the host population.

Moreover, our data showed that the compatible crosses transmitting *Wolbachia* to the next generation had higher reproductive fitness compared to the others. The rates of egg hatching and progeny emergence in the infected female-uninfected male cross were significantly higher than those in infected female-infected male cross. In addition, the number of resultant females from infected female-uninfected male cross was similar to those produced from infected female—infected male cross. These data suggest that the presence of *Wolbachia* in the males have no significant effect on fecundity, fertility and number of female progeny when female is also infected with the same *Wolbachia*.Therefore, in the compatible crosses transmitting *Wolbachia*, female infection matters regardless of male infection.

Considering the reproductive fitness that *Wolbachia* impose on *H. hebetor*, we questioned whether that affects mate choice of the female wasp. Our data showed that infected females chose infected males more frequently than uninfected males. However, there was no difference in male mate choice of the uninfected females. These data suggest that *Wolbachia* infection may modify the mate choice of this wasp. In congruence to these results, it has been reported that in the two-spotted mite, *Tetranychus urticae, *Wolbachia*-infected* females mostly choose infected males to decrease the chance of CI occurring (Vala et al., 2004). Also, it has been shown that *Wolbachia* affect reproduction preference and reproduction isolation among different populations of *D. melanogaster*. This observation may strengthen the hypothesis that infected females are more attractive to males than their uninfected counterparts. This active bias could be due to differences in pheromone compounds of infected and uninfected females that attracts the infected males more than the uninfected males.

## 5. Conclusion

In this study we showed that *Wolbachia* were persistently present during development of the parasitoid wasp, *H. hebetor* highlighting efficient vertical transmission of *Wolbachia*. Different densities of *Wolbachia* during development of *H. hebetor* may be due to specific regulatory mechanisms (such as metamorphosis) that allow it to populate the host at certain stages. By elimination of *Wolbachia* from *H. hebetor* and conducting crossing experiments, we confirmed that *Wolbachia* induced CI in this wasp is most probably through killing female embryos in incompatible crosses. Moreover, we reported that *Wolbachia* affects mate choice and fecundity of *H. hebetor* (in compatible crosses); thereby incur direct fitness to the host that transmits it to the next generation. This strategy facilitates *Wolbachia* spread through populations of the host insects.

## Acknowledgements

Works in Our laboratory (M.M.) is supported by grants from Tarbiat Modares University, National Science Foundation (INSF) and Center for International Scientific Studies and collaboration (CISSC). The authors declare that they have no conflicts of interest with the contents of this article.

## Authors’ contributions

M.M. and Z.B. conceived and designed the experiments. Z.B. performed the experiments. Z.B. and M.M. and A.A.T. analyzed the data and prepared the figures and tables. Z.B., S.A. and M.M. wrote the manuscript. All of the authors reviewed the manuscript.

---

**Fig. S1.**
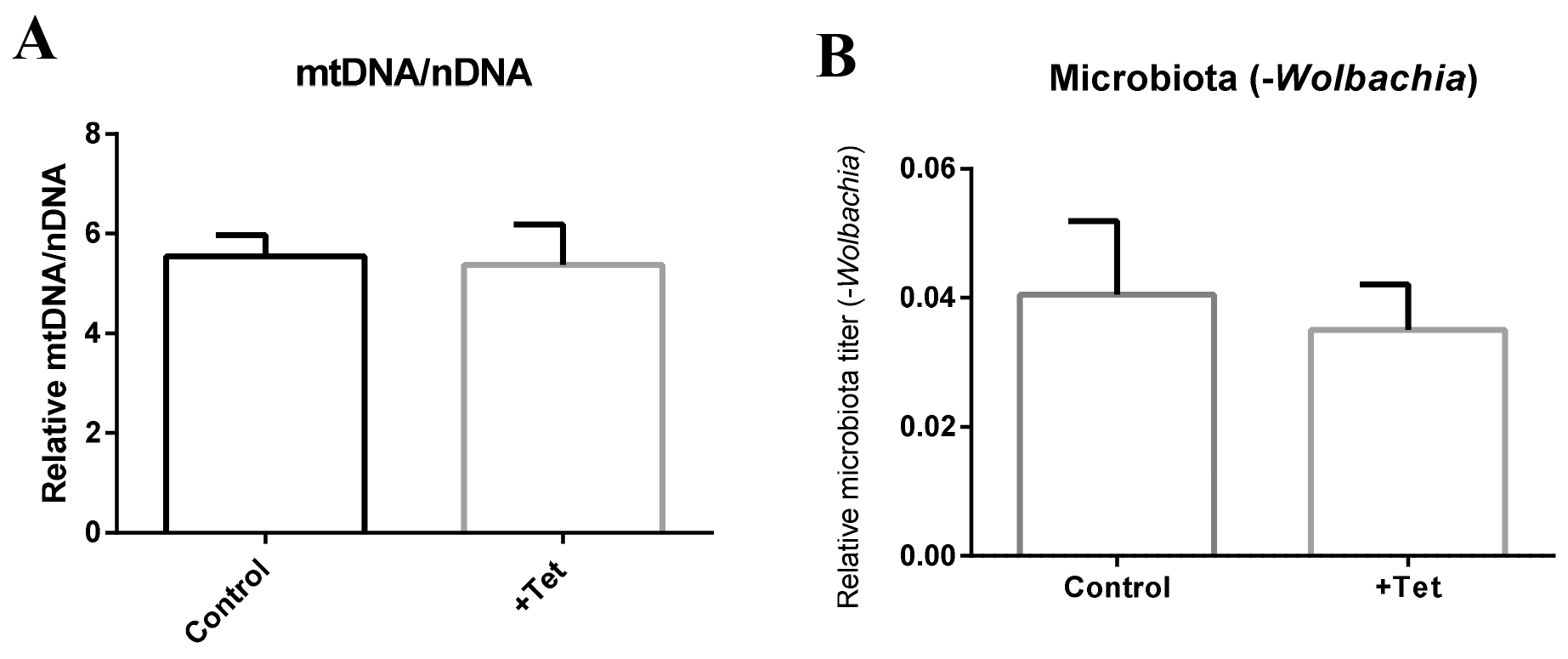
Effect of tetracycline treatment on (A) mitochondria density and (B) microbiota population of *H. hebetor* (8^th^ generation after treatment). There was no significant differences between the control and treated wasps.

## References

Ali, H., et al., 2018. Infection Density Dynamics and Phylogeny of *Wolbachia* Associated with Coconut Hispine Beetle, *Brontispa longissima* (Gestro)(Coleoptera: Chrysomelidae), by Multilocus Sequence Type (MLST) Genotyping. Journal of Microbiology and Biotechnology. 28, 796–808.

Ballard, J., Melvin, R., 2007. Tetracycline treatment influences mitochondrial metabolism and mtDNA density two generations after treatment in *Drosophila*. Insect molecular biology. 16, 799–802.

Beckmann, J. F., et al., 2017. A *Wolbachia* deubiquitylating enzyme induces cytoplasmic incompatibility. Nature Microbiology. 2, 17007.

Bordenstein, S. R., et al., 2003. Host genotype determines cytoplasmic incompatibility type in the haplodiploid genus *Nasonia*. Genetics. 164, 223–233.

Bourtzis, K., et al., 1996. *Wolbachia* infection and cytoplasmic incompatibility in *Drosophila* species. Genetics. 144, 1063–1073.

Breeuwer, J., Werren, J. H., 1993. Cytoplasmic incompatibility and bacterial density in *Nasonia vitripennis*. Genetics. 135, 565–574.

Brownlie, J. C., et al., 2009. Evidence for metabolic provisioning by a common invertebrate endosymbiont, *Wolbachia pipientis*, during periods of nutritional stress. PLoS pathogens. 5, e1000368.

Dedeine, F., et al., 2005. *Wolbachia* requirement for oogenesis: occurrence within the genus *Asobara* (Hymenoptera, Braconidae) and evidence for intraspecific variation in A. tabida. Heredity. 95, 394–400.

Dobson, S., et al., 2004. Fitness advantage and cytoplasmic incompatibility in *Wolbachia* single-and superinfected *Aedes albopictus*. Heredity. 93, 135–142.

Dossi, F. C. A., et al., 2014. Population dynamics and growth rates of endosymbionts during *Diaphorina citri* (Hemiptera, Liviidae) ontogeny. Microbial ecology. 68, 881–889.

Fry, A., et al., 2004. Variable fitness effects of *Wolbachia* infection in *Drosophila melanogaster*. Heredity. 93, 379–389.

Ghimire, M. N., 2008. Reproductive performance of the parasitoid *Bracon hebetor* Say (Hymenoptera: Braconidae) on various host species of Lepidoptera. Oklahoma State University.

Goto, S., et al., 2006. Asymmetrical interactions between *Wolbachia* and Spiroplasma endosymbionts coexisting in the same insect host. Applied and environmental microbiology. 72, 4805–4810.

Hedges, L. M., et al., 2008. *Wolbachia* and virus protection in insects. Science. 322, 702–702.

Hosokawa, T., et al., 2010. *Wolbachia* as a bacteriocyte-associated nutritional mutualist. Proceedings of the National Academy of Sciences. 107, 769–774.

Hunter, M. S., et al., 2003. A bacterial symbiont in the Bacteroidetes induces cytoplasmic incompatibility in the parasitoid wasp *Encarsia pergandiella*. Proceedings of the Royal Society of London B: Biological Sciences. 270, 21852190.

Hurst, G. D., Frost, C. L., 2015. Reproductive parasitism: maternally inherited symbionts in a biparental world. Cold Spring Harbor perspectives in biology. 7, a017699.

Hurst, G. D., et al., 1999. Male-killing *Wolbachia* in two species of insect. Proceedings of the Royal Society of London B: Biological Sciences. 266, 735–740.

Jiggins, F. M., et al., 1998. Sex ratio distortion in *Acraea encedon* (Lepidoptera: Nymphalidae) is caused by a male-killing bacterium. Heredity. 81, 87–91.

Kageyama, D., et al., 2010. Detection and identification of *Wolbachia* endosymbionts from laboratory stocks of stored-product insect pests and their parasitoids. Journal of stored products research. 46, 13–19.

Karamipour, N., et al., 2016. Gammaproteobacteria as essential primary symbionts in the striped shield bug, *Graphosoma lineatum* (Hemiptera: Pentatomidae). Scientific reports. 6, 33168.

Koukou, K., et al., 2006. Influence of antibiotic treatment and *Wolbachia* curing on sexual isolation among *Drosophila melanogaster* cage populations. Evolution. 60, 87–96.

Kruse, A., et al., 2017. Combining’omics and microscopy to visualize interactions between the Asian citrus psyllid vector and the Huanglongbing pathogen Candidatus Liberibacter asiaticus in the insect gut. PloS one. 12, e0179531.

Laven, H., 1967. Eradication of *Culex pipiens* fatigans through cytoplasmic incompatibility. Nature. 216, 383–384.

LePage, D., Bordenstein, S. R., 2013. *Wolbachia*:. can we save lives with a great pandemic? Trends in parasitology. 29, 385–393.

Miller, W. J., et al., 2010. Infectious speciation revisited: impact of symbiont-depletion on female fitness and mating behavior of Drosophila paulistorum. PLoS Pathogens. 6, e1001214.

Ming, Q.-L., et al., 2015. *Wolbachia* infection dynamics in *Tribolium confusum* (Coleoptera: Tenebrionidae) and their effects on host mating behavior and reproduction. Journal of economic entomology. 108, 1408–1415.

Narita, S., et al., 2007. Naturally occurring single and double infection with *Wolbachia* strains in the butterfly *Eurema hecabe*: transmission efficiencies and population density dynamics of each *Wolbachia* strain. FEMS microbiology ecology. 61, 235–245.

Noda, H., et al., 2001. Infection density of *Wolbachia* and incompatibility level in two planthopper species, *Laodelphax striatellus* and *Sogatella furcifera*. Insect biochemistry and molecular biology. 31, 727–737.

O’Neill, S. L., et al., 1992. 16S rRNA phylogenetic analysis of the bacterial endosymbionts associated with cytoplasmic incompatibility in insects. Proceedings of the National Academy of Sciences. 89, 2699–2702.

O’Neill, S. L., Karr, T. L., 1990. Bidirectional incompatibility between conspecific populations of *Drosophila simulans*. Nature. 348, 178.

Okayama, K., et al., 2016. Costs and benefits of symbiosis between a bean beetle and *Wolbachia*. Animal Behaviour. 119, 19–26.

Pan, X., et al., 2017. The bacterium *Wolbachia* exploits host innate immunity to establish a symbiotic relationship with the dengue vector mosquito *Aedes aegypti*. The ISME journal. 12, 277.

Peng, Y., et al., 2008. *Wolbachia* infection alters olfactory-cued locomotion in *Drosophila* spp. Applied and environmental microbiology. 74, 3943–3948.

Perrot-Minnot, M.-J., et al., 1996a. Single and double infections with *Wolbachia* in the parasitic wasp *Nasonia vitripennis* effects on compatibility. Genetics. 143, 961972.

Perrot-Minnot, M., et al., 1996b. Double and single *Wolbachia* infections in *Nasonia vitripennis*. Genetics. 143, 961–972.

Rohrscheib, C. E., et al., 2015. *Wolbachia* influences the production of octopamine and affects Drosophila male aggression. Applied and environmental microbiology. 81, 4573–4580.

Rousset, F., et al., 1992. *Wolbachia* endosymbionts responsible for various alterations of sexuality in arthropods. Proceedings of the Royal Society of London B: Biological Sciences. 250, 91–98.

Serbus, L. R., et al., 2008. The genetics and cell biology of *Wolbachia*-host interactions. Annual review of genetics. 42, 683–707.

Sinkins, S., et al., 1995. *Wolbachia pipientis*: bacterial density and unidirectional cytoplasmic incompatibility between infected populations of Aedes albopictus. Experimental parasitology. 81, 284–291.

Stouthamer, R., et al., 1990. Antibiotics cause parthenogenetic *Trichogramma* (Hymenoptera/Trichogrammatidae) to revert to sex. Proceedings of the National Academy of Sciences. 87, 2424–2427.

Stouthamer, R., Mak, F., 2002. Influence of antibiotics on the offspring production of the *Wolbachia*-infected parthenogenetic parasitoid Encarsia formosa. Journal of invertebrate pathology. 80, 41–45.

Stouthamer, R., Werren, J. H., 1993. Microbes associated with parthenogenesis in wasps of the genus *Trichogramma*. Journal of Invertebrate Pathology. 61, 6–9.

Tagami, Y., et al., 2006. *Wolbachia*-induced cytoplasmic incompatibility in *Liriomyza trifolii* and its possible use as a tool in insect pest control. Biological Control. 38, 205–209.

Tortosa, P., et al., 2010.*Wolbachia* age-sex-specific density in *Aedes albopictus*: a host evolutionary response to cytoplasmic incompatibility? PLoS One. 5, e9700.

Vala, F., et al., 2004. *Wolbachia* affects oviposition and mating behaviour of its spider mite host. Journal of evolutionary biology. 17, 692–700.

Vasquez, C. J., et al., 2011. Discovery of a CI-inducing *Wolbachia* and its associated fitness costs in the biological control *agent Aphytis* melinus DeBach (Hymenoptera: Aphelinidae). Biological Control. 58, 192–198.

Vavre, F., et al., 2000. Evidence for female mortality in *Wolbachia*-mediated cytoplasmic incompatibility in haplodiploid insects: epidemiologic and evolutionary consequences. Evolution. 54, 191–200.

Vavre, F., et al., 2002. Infection polymorphism and cytoplasmic incompatibility in Hymenoptera-*Wolbachia* associations. Heredity. 88, 361.

Vavre, F., et al., 1999. Phylogenetic status of a fecundity-enhancing *Wolbachia* that does not induce thelytoky in *Trichogramma*. Insect molecular biology. 8, 67–72.

Wang, X.-X., et al., 2017. Incomplete removal of *Wolbachia* with tetracycline has two-edged reproductive effects in the thelytokous wasp Encarsia formosa (Hymenoptera: Aphelinidae). Scientific Reports. 7.

Weinert, L. A., et al., 2015. The incidence of bacterial endosymbionts in terrestrial arthropods. Proc. R. Soc.B. 282, 20150249.

Werren, J. H., et al., 2008. *Wolbachia*:. master manipulators of invertebrate biology. Nature Reviews Microbiology. 6, 741.

Yen, J. H., Barr, A. R., 1971. New hypothesis of the cause of cytoplasmic incompatibility in Culex pipiens L. Nature. 232, 657–658.

Yixin, H. Y., et al., 2013. *Wolbachia*-associated bacterial protection in the mosquito *Aedes aegypti*. PLoS neglected tropical diseases. 7, e2362.

Zchori-Fein, E., et al., 1992. Male production induced by antibiotic treatment in Encarsia formosa (Hymenoptera: Aphelinidae), an asexual species. Experientia. 48, 102105.

Zhao, D.-X., et al., 2013. *Wolbachia*-host interactions: host mating patterns affect *Wolbachia* density dynamics. PLoS One. 8, e66373.

